# Molecular genetic characterization of *CASEIN KINASE 1-LIKE 12* in *Arabidopsis*

**DOI:** 10.1101/2025.09.17.676859

**Authors:** Adam Seluzicki, Annemarie E. Branks, Sowmya Poosapati, Joanne Chory

## Abstract

The CASEIN KINASE 1 (CK1) family plays diverse roles in development, physiology, and disease in eukaryotes. In *Arabidopsis thaliana* the CASEIN KINASE 1-LIKE (CKL) family has 13 members, but to date the roles of these kinases remain largely unclear. Here we characterize several insertion mutants, finding that CKL12 may contribute to hypocotyl and in primary root growth. Differential effects of insertions in different parts of the gene suggest that the 3’ end of the transcript may be important for CKL12 function. We provide evidence that *CKL12* may be a transcriptional target of brassinosteroid (BR) signaling. The *CKL12* promoter contains in-vitro binding sites for BR-related transcription factors. Knock-down of these transcription factors using RNA interference reduces *CKL12* transcript. Together, these data suggest that CKL12 may act downstream of BR signaling to regulate seedling growth.

## INTRODUCTION

The CASEIN KINASE 1 (CK1) family of serine/threonine kinases is well described in eukaryotes, playing central signaling roles in controlling the circadian clock and regulating cellular signaling in development^1–4^. CK1 proteins have highly conserved compact kinase domains, followed by highly variable C-terminal regulatory domains with auto-inhibitory properties^5–7^. The *Arabidopsis* genome contains 13 orthologs of CK1, with closest homology to the CK1δ/ε group described in animals. The *Arabidopsis* CKL family contains three subgroups (CKL-A, -B, -C), and it appears that several members have undergone duplications^8^. Four additional genes (MUT9-LIKE KINASES (MLKs) or PHOTOREGULATORY PROTEIN KINASES (PPKs)), show shared homology with the CK1 kinase domain but have larger C-terminal extensions^9^.

Despite the clear importance of CK1 kinases in many systems, there are relatively few studies examining them in plants. To date most members of the CKL family remain uncharacterized. Pharmacological studies have shown that broad inhibition of CASEIN KINASE 1-LIKE (CKL) proteins in *Arabidopsis* results in altered progression of the circadian clock, possibly via through effects on PRR5 and TOC1^8–10^. Similar results were obtained with broad knock-down of the CKL family using RNA interference, although no single CKL was found to be uniquely important for these effects^10^. CKL3 and CKL4 were implicated in control of blue light signaling via direct phosphorylation of CRYPTOCHROME 2, although a later study mapped this activity to the PPK subgroup^11,12^. CKL6 was shown to associate with microtubules through the C-terminal domain, directly phosphorylating TUBULIN beta 3, as well as localizing to late endosomal vesicles^13,14^. CKL2 was described as a positive regulator of Abscisic Acid (ABA) responses, including germination, root growth, and proline accumulation^15^. CKL2 was also found to be a factor regulating crosstalk between the ABA and Brassinosteroid (BR) hormone signaling pathways, being activated by ABA during stress to directly phosphorylate the BR receptor BRI1, priming it for rapid recovery after stress has passed^16^. There is little information regarding CKL12 in *Arabidopsis*. One study generated promoter-reporter constructs to examine expression patterns of the CKL family in different plant tissues, finding that the CKL12 promoter could drive expression in the vasculature, trichomes, and anthers, as well as the root cap, root tip, and primary root^17^.

Brassinosteroid hormone signaling is an central pathway controlling plant growth. Loss of function in this pathway results in dwarf plants that show a photomorphogenic phenotype when grown in dark. The hypocotyl fails to elongate, and the apical hook and cotyledons open similar to light-grown plants^18–23^. Genetic screens for suppressors of these phenotypes uncovered dominant mutations in the BR-responsive transcription factors BRASSINAZOLE RESISTANT 1 (BZR1) and BRI1 EMS SUPPRESSOR 1 (BES1)^24,25^. The *bzr1-1D* and *bes1-1D* mutants both mimic mild BR deficiency in long-day (LD) conditions, and rescue BR deficiency in constant dark (DD). The search for additional homologs of these TFs revealed a small family of BES1/BZR1 HOMOLOG (BEH) genes that also contribute to transcription downstream of BR signaling^26^. The downstream transcription factors in the BR pathway also have feedback effects on BR hormone synthesis, and interact with other key signaling pathways including auxin and gibberellin, as well as light signaling^25,27–31^.

Here we identify genetic reagents, characterize seedling growth, and analyze transcriptional regulation of *CASEIN KINASE 1-LIKE 12* (*CKL12*). We find that insertion mutations disrupting the 3’ end of the gene may cause mild reductions in growth in both hypocotyl and primary root, while a possible null mutant has weaker effects. We also find that BR-related transcription factors are likely to play a role in promoting *CKL12* transcription. Together, this study provides initial characterization of a member of the under-explored CKL gene family in *Arabidopsis*.

## MATERIALS and METHODS

### Plant Material

*Arabidopsis thaliana* plants were propagated on soil with added fertilizer and fungicide. Insertion mutants for *CKL12* (AT5G57015) were obtained from the Arabidopsis Biological Resource Center (ABRC - The Ohio State University). Insertion lines were confirmed by PCR from genomic DNA using primers in Supplementary Table 1. Lines carrying *bzr1-1D* and *bes1-1D* were described^24,25^. RNAi lines targeting *BES1*/*BZR1* were described^26^.

### Growth Conditions

Seeds were surface-sterilized using chlorine gas and plated on 1/2x Linsmaier & Skoog medium + 0.8% phytoagar (Caisson). Plates were stratified in the dark at 4°C for 4 days. Plates were then moved to growth chambers (Percival) under appropriate conditions: LD (16h light-8h dark, light intensity: 100 μmol m^−2^s^−1^, 20°C), LL (constant light, 100 μmol m^−2^s^−1^, 20°C), DD (constant darkness, 20°C). Light source was cool-white fluorescent bulbs (Philips). Seeds to be grown under DD were exposed to white light to trigger germination, then wrapped in foil and grown in the same chamber as LD or LL plates. Plates were scanned and measured using ImageJ software (NIH)^32^.

### qRT-PCR

RNA was extracted from seedlings collected at the indicated time points using the RNeasy Plant Mini Kit (Qiagen) and treated with DNAse according to manufacturer’s instructions. Maxima First Strand cDNA Synthesis Kit was used to make cDNA from 2μg RNA from each sample. Reactions were run using SYBR-Green detection in a BioRad CFX Opus 384 Real-Time PCR Machine. IPP2 (AT3G02780) transcript was used as the internal control for quantification using the ΔΔCq method. Semi-quantitative RT-PCR was done using samples previously used for qRT-PCR, run for 40 PCR cycles (annealing: 56°C for 30s, elongation: 72°C for 45s) and run on 2% agarose gel with ethidium bromide visualization.

### Statistics

Student’s T-Test (two-tailed) was used for comparison between two samples. One-way ANOVA + Tukey’s HSD test (α=0.05) was used to compare three or more samples.

## RESULTS

### Protein sequence alignment of CKL12 with human and fly CK1ε

Examination of the protein sequence of CKL12 in relation to the human and *Drosophila* CK1ε shows that the key residues in the function of the kinase domain are conserved across evolutionary time. The amino acid residues that form the ATP-binding pocket, the active site, and the location of mutations identified in mammals (human, hamster) and in *Drosophila* are indicated on the alignment^33,34^. The ATP-binding pocket (orange) and active site residues (magenta) of CKL12 are identical to those of HsCK1 and DmCK1 (Figure 1). Proline 47, which is mutated to serine in the *Drosophila dbt*^*S*^ mutant, is conserved. However, methionine 80, which is mutated to isoleucine in *dbt*^*L*^, is leucine in CKL12, suggesting that the kinase of CKL12 may be reduced relative to other family members^35^. All three residues that form the anion-binding motif, one of which is notable for being mutated in the *tau* mutant hamster (R178C) are retained in CKL12^34^. The C-terminal regions of all three proteins are of similar length, but contain unique sequence.

**FIGURE 1.**
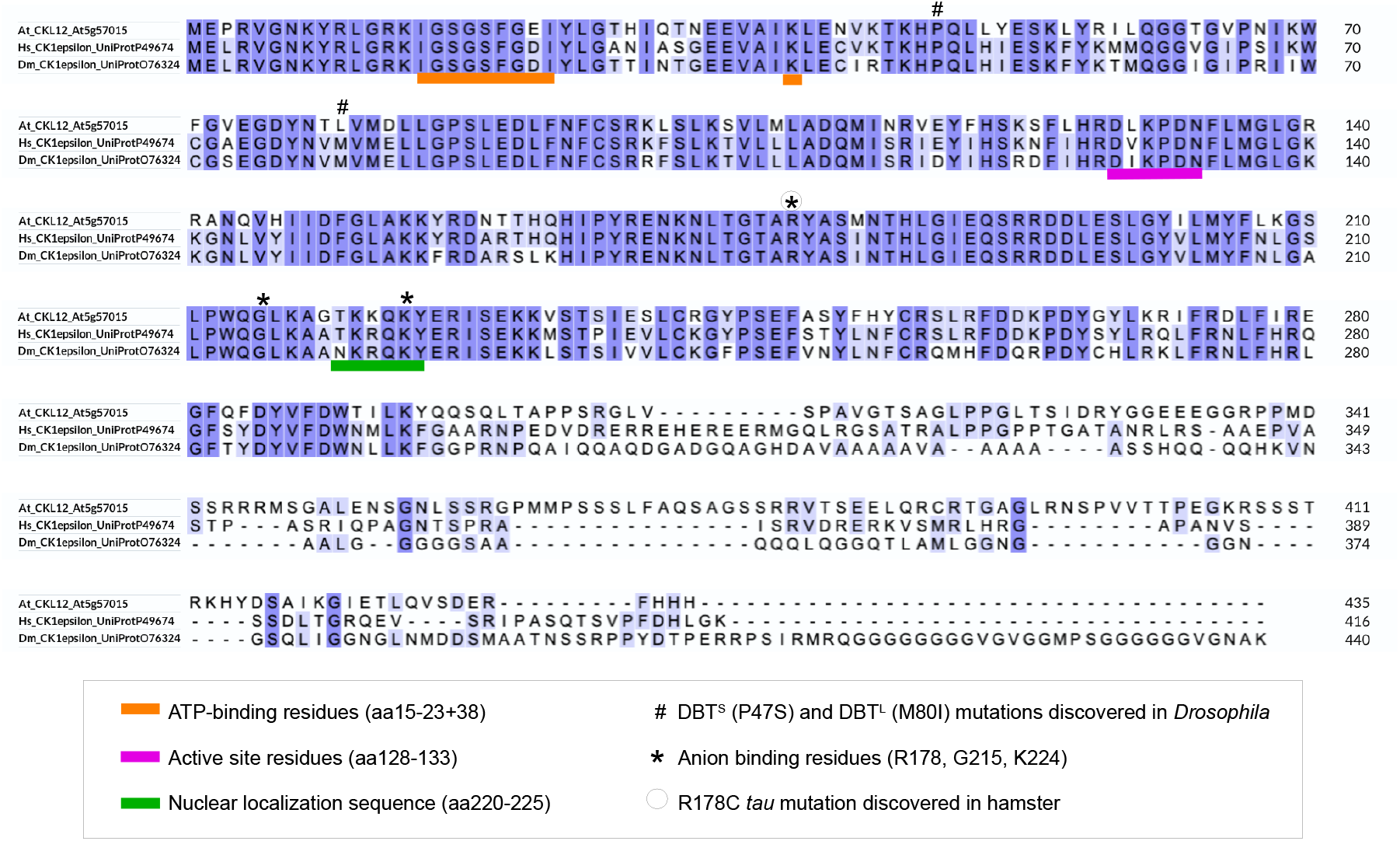
Protein sequence alignment of CKL12 with human and *Drosophila* CK1ε. Sequence alignment is annotated with features identified in mammalian and Drosophila CK1. Features include the ATP-binding domain (Orange underscore); Active site (Magenta underscore); Nuclear Localization Sequence (Green Underscore); Amino acids altered in Drosophila DBTS and DBTL mutants (#); Anion binding site residues (*). One of the anion-binding residues, R178, is mutated in the *tau* mutant hamster and is indicated by a circled asterisk.

### Isolation and molecular characterization of T-DNA insertion alleles of CKL12

In the effort to expand the set of reagents available to study the CKL family in *Arabidopsis*, we isolated and confirmed three insertion alleles of CKL12. These three insertions are in three distinct regions of the gene, with different expected consequences. SALK_012002 (referred to from here on as *ckl12-1*) is inserted in the 3’ untranslated region (UTR), SALK_059455 (*ckl12-2*) is inserted in the 12th intron, and SALK_119322 (*ckl12-3*) is inserted in the second intron (Figure 2A). The insertion site and zygosity of each of these insertions was confirmed by PCR, and lines were propagated from homozygous individuals (Fig. 2B-D).

**FIGURE 2.**
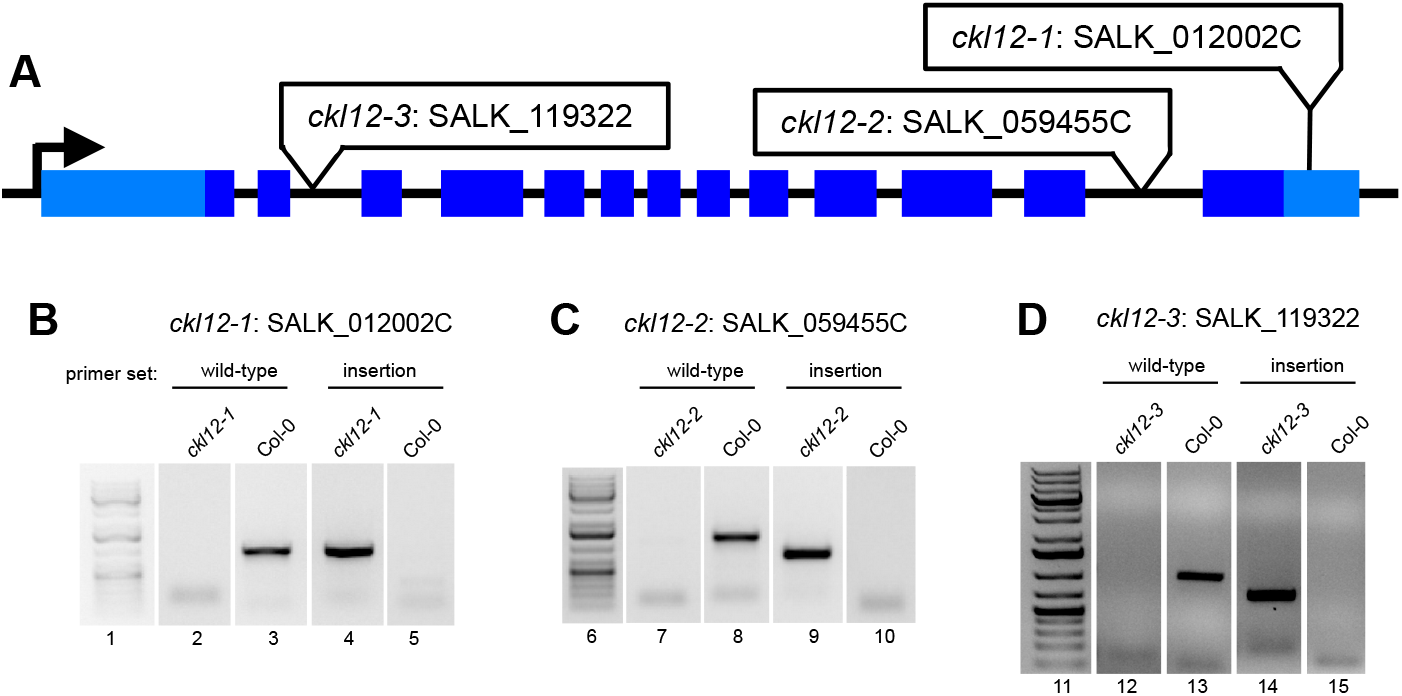
Isolation and confirmation of TDNA insertion mutants in CKL12. (A) Structure of the CKL12 gene locus. 5’ and 3’ untranslated regions are light blue rectangles. Exons are in dark blue rectangles. Insertion sites for each allele are indicated in white boxes. (B-D) PCR amplified products from genomic DNA isloated from wild-type or mutant plants. Primer sets targeting wild-type genomic DNA flank the TDNA insertion site. Primers targeting the TDNA alleles use one primer within the TDNA and one of the primers used for wild-type amplification. Numbers below the gel images indicate the region of the unspliced images marked in Supplemental Figure 1.

To examine the molecular consequences of each insertion on the production of *CKL12* transcript, we carried out quantitative reverse-transcription polymerase chain reaction (qRT-PCR) assays. Primer sets used are diagrammed in Fig. 3A. Analysis of *ckl12-2*, inserted in intron 12, showed wild-type transcript levels when assayed with primers across exons 5-6, but failed to amplify across exons 12-13, suggesting that the T-DNA insertion disrupts the 3’ end of the coding region (Fig. 3B). CKL12 transcript level assayed with the “mid” primer set was similar to wild-type in *ckl12-1* but was significantly elevated in *ckl12-3* (Figure 3C). Analysis of the same cDNA from Fig. 3C using the 5’ primer set, which surrounds the *ckl12-3* insertion site, amplified from the Col-0 and *ckl12-1* samples, but did not amplify from *ckl12-3* samples (Fig.3D). Thus, *ckl12-3* could potentially result in over-expression of a CKL12 fragment that lacks the first 25 amino acids encoding the ATP-binding pocket (Fig. 1). Together, these data indicate *ckl12-3* is a likely loss-of-function allele, while *ckl12-1* and *ckl12-2* may modify CKL12 protein function in more complex ways.

**FIGURE 3.**
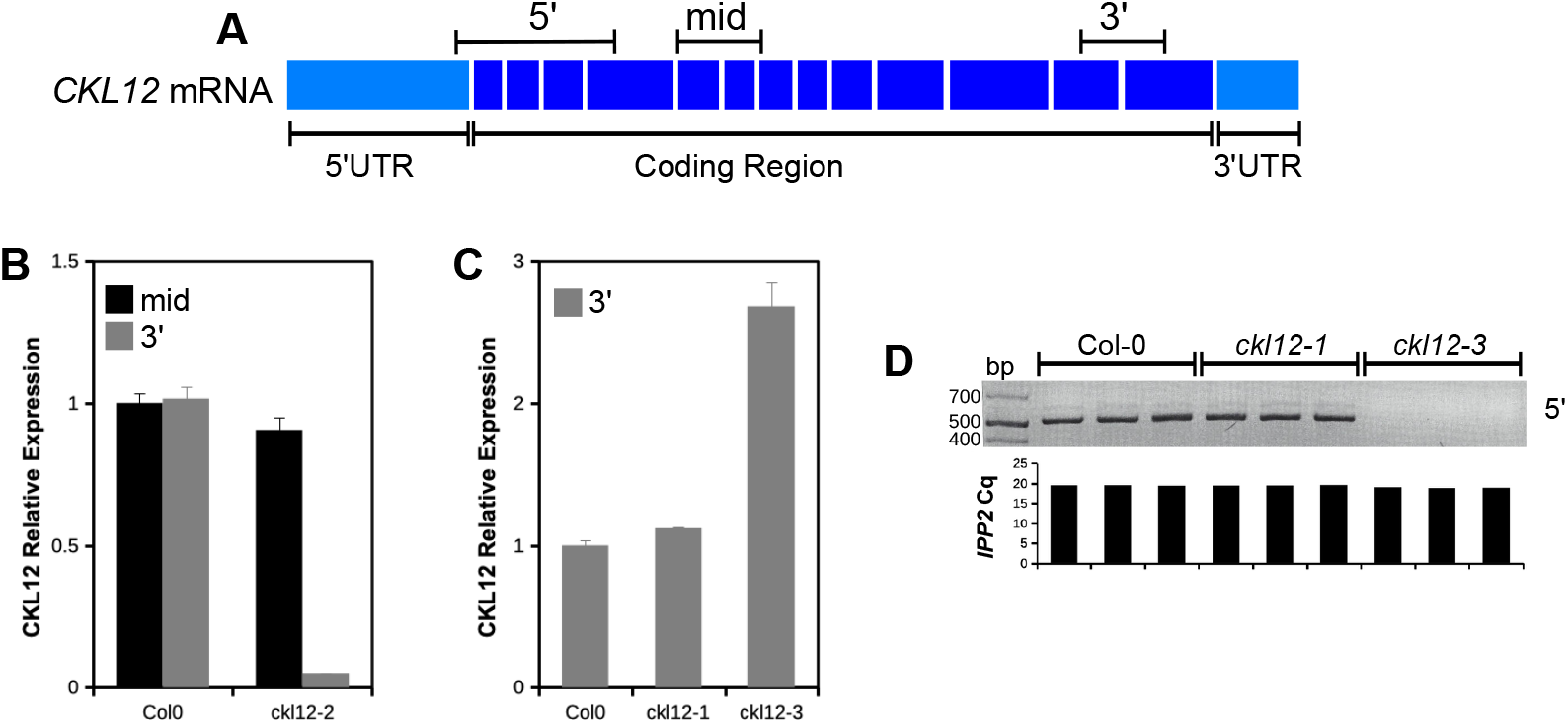
Analysis of *CKL12* transcript in TDNA mutant plants. (A) Model of mature *CKL12* mRNA. Target amplicons are indicated above the model. (B,C) qRT-PCR of *CKL12* transcript. Transcript abundance was quantified using the ΔΔCq method with *IPP2* as the internal control. Experiments were done in biological triplicate. (B) *CKL12* transcript in *ckl12-2* using the mid (black bars) and 3’ (gray bars) amplicons sampled on day 6. (C) qRT-PCR of *ckl12-1* and *ckl12-3* mutants using the 3’ (gray bars) amplicon sampled on day 4. (D) Semi-quantitative RT-PCR of *CKL12* transcript using the 5’ amplicon in the same cDNA samples as (C). Cq values for *IPP2* in each replicate from (C) are aligned to each sample beneath the gel image.

### CKL12 may promote growth in the hypocotyl and root

To begin to understand the function of CKL12, we tested seedling growth in *ckl12* mutants. Under long day photoperiod (LD16:8) at 20°C, *ckl12-2* and *ckl12-3* showed slight but statistically shorter hypocotyls than control, while *ckl12-1* was comparable to control, all of which were ~95% of Col-0 (Fig. 4A,B,G,H). Dark (DD) grown *ckl12-1* and *ckl12-2* seedling hypocotyls were shorter than control, showing ~10% reductions in both hypocotyl and primary root lengths (Fig. 4C,I). Primary roots were shorter than control in *ckl12-1* and *ckl12-2* (Fig. 4D,G), but comparable to control in *ckl12-3* (Fig. 4E,H). The short root phenotype in *ckl12-1* and *ckl12-2* was maintained in DD (Fig. 4F,I). Together, these data suggest that disruption, but not elimination, of CKL12 may disrupt growth in both hypocotyl and root.

**FIGURE 4.**
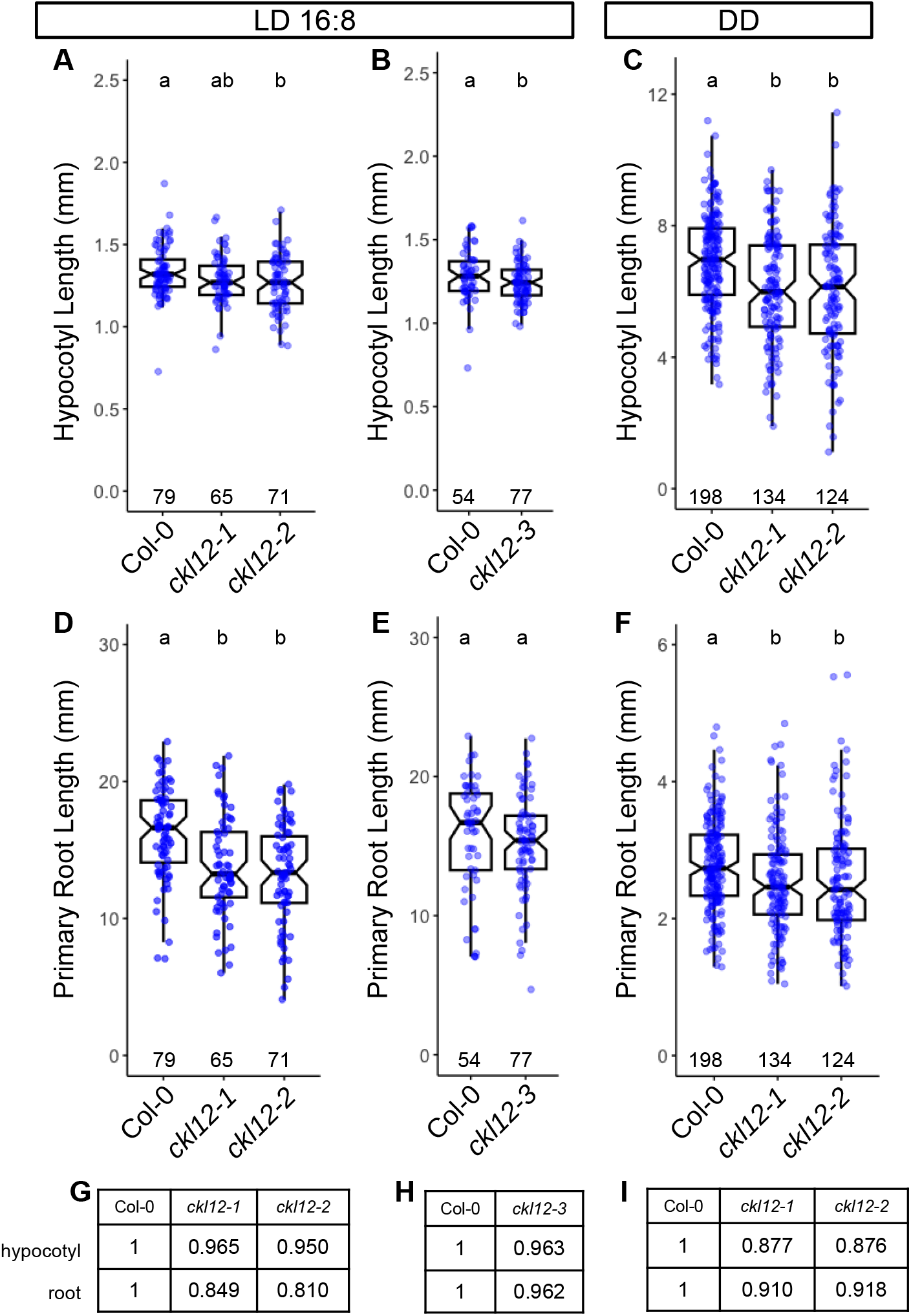
Hypocotyl and root growth in *ckl12* mutants. (A-C) Hypocotyl length. (D-F) Primary root length. (A,B,D,E) Seedlings grown under LD16:8, measured on day 6. (C,F) Seedlings grown in constant dark (DD), measured on day 3. Box plots show pooled data from independent replicate experiments [(A,D) 3, (B,E) 2, (C,F) 5]. One experiment included *ckl12-1, ckl12-2*, and *ckl12-3*. Col-0 data from this experiment are included in both (A,B) and (D,E) for hypocotyl and root measurements, respectively. Mean relative Hypocotyl and Primary Root lengths in LD (G,H) and DD (I), normalized to Col-0=1, are noted in tables. Box plots indicate the median, 25th and 75th percentiles. Whiskers extend to 1.5*interquartile range. Notches approximate the 95% confidence interval of the median. N (individual plants scored) is shown near the x-axis for each genotype. Different letters at top indicate statistically significant difference between groups (α=0.05) by one-way ANOVA+Tukey HSD.

### CKL12 expression is regulated by brassinosteriod-related transcription factors

We hypothesized that transcriptional regulation of CKL12 may provide clues as to it’s function. We searched the DAP-seq database for transcription factor binding sites in the promoter region of *CKL12*, comprised of ~600bp upstream of the transcription start site to the nearest neighboring gene. We found annotated binding sites for BZR1 and BES1, two key transcription factors of the brassinoteroid (BR) signaling pathway (Fig. 5A)^36^. We hypothesized that the BR pathway may regulate *CKL12* expression. We assayed *CKL12* transcript in several lines that have modified BR TFs in 10-day-old seedlings grown in LD conditions. The *bzr1-1D* and *bes1-1D* mutants both expressed *CKL12* at levels comparable to control. Two *bes1-RNAi* lines, which knock down both *BES1* and *BZR1*, reduce *CKL12* transcript (Fig. 5B)^26^. The stronger of the two lines, 14-08i, reduces *CKL12* expression by approximately half.

**FIGURE 5.**
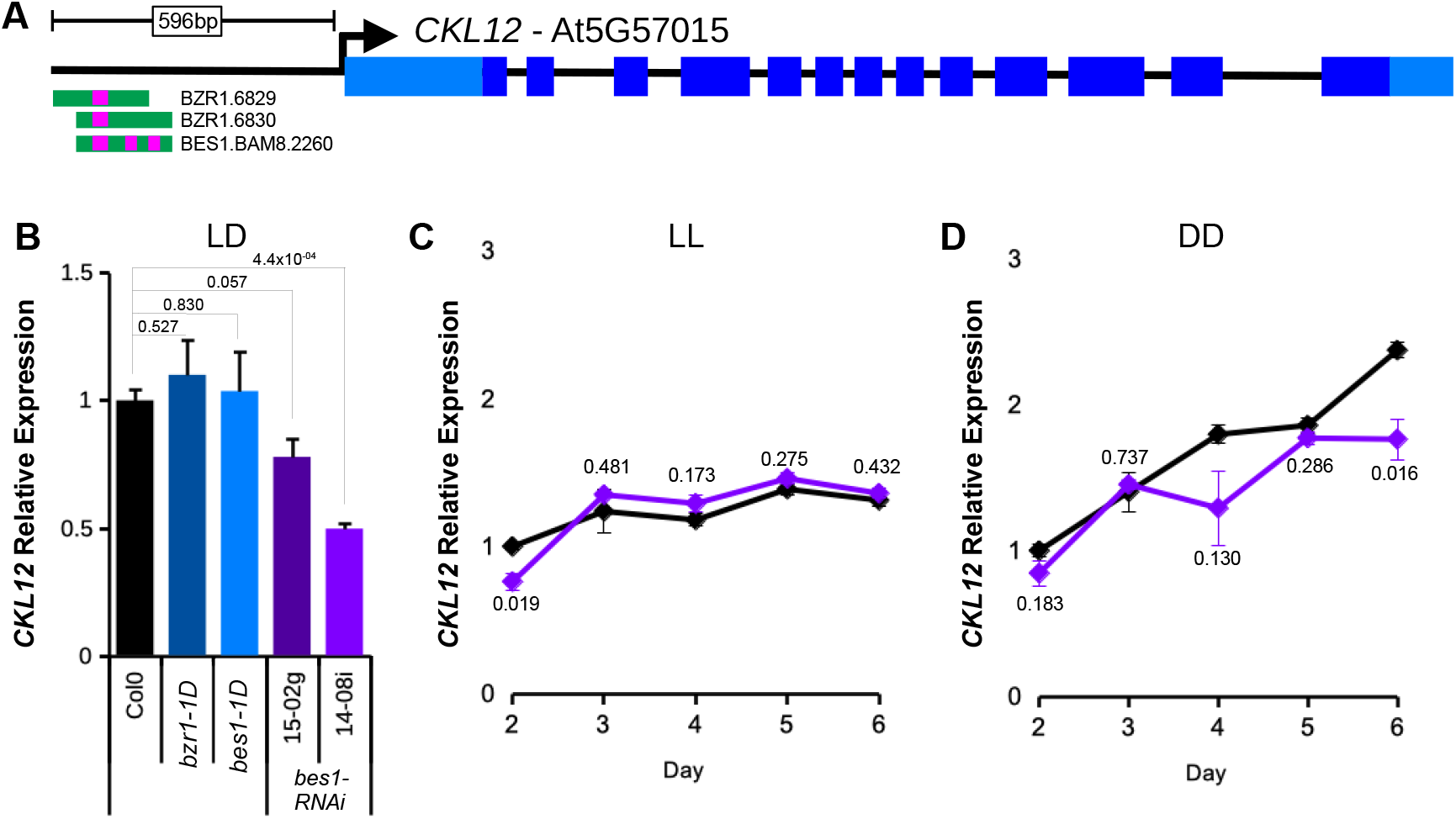
*CKL12* transcription may be promoted by brassinosteroid-related transcription factors in darkness. (A) Diagram derived from the DAP-seq browser track for the *CKL12* locus. *CKL12* gene structure is shown at top. The *CKL12* promoter (from the *CKL12* transcription start site up the the 3’ end of the upstream gene transcript) is 596bp. Transcription factor binding sites are shown below the gene model, with BES1 and BZR1 binding sites noted with magenta bars. (B) qRT-PCR of *CKL12* in 10-day old seedlings grown in LD16:8. Genotypes: *bzr1-1D* (dark blue), *bes1-1D* (light blue), and *bes1-RNAi* lines 15-02g (dark purple) and 14-08i (light purple). Transcript abundance was determined using the ΔΔCq method, using *IPP2* as the internal control in biological triplicate. Error bars are SEM. Two-tailed T-test p-values for each mutant vs. Col-0 are indicated above the bars. (C,D) Time course expression of CKL12 in Col-0 (black) and *bes1-RNAi* 14-08i (light purple) on days 2-6 grown in LL (C) and DD (D). Expression level was determined using the ΔΔCq method using *IPP2* as the internal control, normalized to Col-0 on day 2, within each light condition. Samples in B-D were assayed using the 3’ primer set diagramed in Fig. 3. Samples are in biological triplicate with the exception of *bes1-RNAi* in DD on day 4, which is in duplicate. Error bars are SEM. Two-tailed T-test p-values for *bes1-RNAi* vs. Col-0 are included near the traces.

BR signaling is required for hypocotyl growth in the dark. We observed reduced hypocotyl elongation in the dark in *ckl12* mutants. Thus, we examined CKL12 transcript in constant light (LL) and constant dark (DD) conditions in Col-0 and bes1-RNAi plants across days 2-6. In LL, *CKL12* transcript shows a very mild increase from day 2 to day 6, and expression is similar in Col-0 and *bes1-RNAi (14-08i)* plants, with the exception of lower expression on day 2 in the RNAi line (Fig. 5C). In DD, *CKL12* transcript increases more rapidly, more than doubling from day 2 to day 6 in Col0 (Fig. 5D). The *bes1-RNAi* plants show attenuated CKL12 transcript accumulation, with the strongest effect on day six. Together, these data support the hypothesis that BR-related transcription factors may be important transcriptional regulators of *CKL12*-expression.

## DISCUSSION

CK1 proteins are important regulators of many signaling processes in eukaryotes, but are little studied in plants. In this study, we have provided genetic and molecular characterization of *CKL12*, a rarely studied member of the CASEIN KINASE 1-LIKE family in *Arabidopsis*. We have isolated and confirmed three insertion mutant alleles, and characterized the consequences of these insertions on gene expression (Figs. 2,3). There are other insertions in the *CKL12* genomic locus, with most mapping to the last intron near the site of *ckl12-2*. It is unclear why this is such a “hot spot” for insertions, but it is possible that there are sequence-intrinsic properties that enhance integration. Indeed, T-DNA insertions are known to preferentially integrate in regions of low GC content, and the last intron of *CKL12* is only 29% GC^37^. We isolated *ckl12-3*, which is inserted in the second intron and strongly disrupts the *CKL12* transcript (Figs. 2D, 3C-D). Interestingly, this allele shows the strongest disruption, and the weakest phenotypes of the alleles we tested (Fig. 4G-I). The pROK2 vector that comprises the insertion vector in *ckl12-3* contains a 35S promoter which is the likely cause of the high expression detected 3’ of the insertion^38^. However, the lack of contiguous transcript in the 5’ end of the gene, connecting the regions encoding the ATP-binding pocket and the catalytic site, suggests that this allele would be unable to produce functional protein (Fig. 3D). These results support the hypothesis that *CKL12* is not an essential gene, and that the uneven distribution of insertion sites is more likely due to sequence preference than negative consequences of disrupting the gene.

Two of the three alleles that we characterized were inserted in the 3’ end of the gene: *ckl12-1* in the 3’ UTR, and *ckl12-2* in intron 12 (Fig. 2A-C). Both insertions had minimal effects on transcript levels (Fig. 3B-C). However, *ckl12-1* was inserted in the 3’UTR, potentially altering translation, localization, or binding of accessory proteins^39^. Additionally, *ckl12-2* disrupted the 3’ end of the transcript, potentially removing the C-terminal 72 amino acids (Figs. 1, 3B). These two alleles showed similar phenotypes. The C-terminal tails of CK1 proteins are often auto-inhibitory. It is possible then, that removing a portion of this tail, as in *ckl12-2*, may reduce auto-inhibition and lead to a more active kinase. Similarly, modification of the 3’UTR, as in *ckl12-1*, may lead to enhanced, or un-regulated, translation. Thus, we may be observing the phenotypic consequences of hyperactivity or over-production of CKL12 in these mutants.

The CKL family in *Arabidopsis* contains 13 members, and sequence analysis has suggested that this family has expanded via whole-genome or other duplications in the recent past^17^. It is likely that these genes retain redundant functions. We find that the three alleles we tested show mild phenotypes, with the strongest two alleles showing ~10% reductions in hypocotyl and root length (Fig. 4). If, as discussed above, the strongest phenotypes arise from overactive CKL12, with minimal consequences in the expected loss-of-function mutant, deeper analysis of CKL functions will likely require both over-expression constructs, including truncations removing the C-terminal tails, and higher-order mutants. *CKL12* fits into a clade with *CKL1, 2*, and *5*, with *CKL1* being its closest paralog^17^. It may therefore be necessary to generate double, triple, or quadruple mutants covering all members of this clade to fully examine the roles of CKL genes in *Arabidopsis*.

Three independent lines of evidence indicate possible brassinosteroid involvement in *CKL12* function. 1) We observed reduced hypocotyl and root growth, both of which are regulated by BR, in *ckl12* mutants. 2) Published data show that the *CKL12* promoter is bound in vitro by the BR-related transcription factors BZR1 and BES1^36^. 3) RNAi knock-down of *BZR1* and *BES1* results in reduced *CKL12* transcript. The BR signaling pathway bears similarity to the WNT/β-catenin pathway in animals, with both being activated by extracellular signals through transmembrane receptors in concert with co-receptor proteins^24^. Both have GSK3β-type kinases as central signal transduction factors. In animals, CK1 and CK2 play important roles in the WNT signaling complex. Specific functions of CKL12 in the canonical BR signaling pathway are unknown at present, although CKL2 has been implicated in cross-regulation of BR signaling by ABA-activated direct phosphorylation of the BR receptor BRI1^16^. BR biosynthesis is feedback-regulated, with expression of BR biosynthetic enzymes and production of BR metabolic intermediates being down-regulated by active BR signaling and in the stabilized *bzr1-1D* mutant^25^. CKL-family kinases, possibly including CKL12 and CKL2, may also be regulated by a feedback mechanism, fine tuning BR signal strength via transcriptional output of BZR1/BES1 transcription factors.

## ACKNOWLEDGEMENTS

This work was carried out under the supervision of Joanne Chory (Investigator, HHMI - deceased) and supported by National Institutes of Health (NIH) grant R35-GM122604 to Joanne Chory, and by the Howard Hughes Medical Institute.

## AUTHOR CONTRIBUTIONS

A.S. and A.E.B. designed and carried out research and analyzed data. S.P. carried out research. A.S. wrote the paper with input from A.E.B. and S.P. J.C. designed and supervised research, and acquired funding.

## DATA AVAILABILITY

Data supporting the conclusions of this paper are included in the figures.

---

**Supplementary Figure 1.**
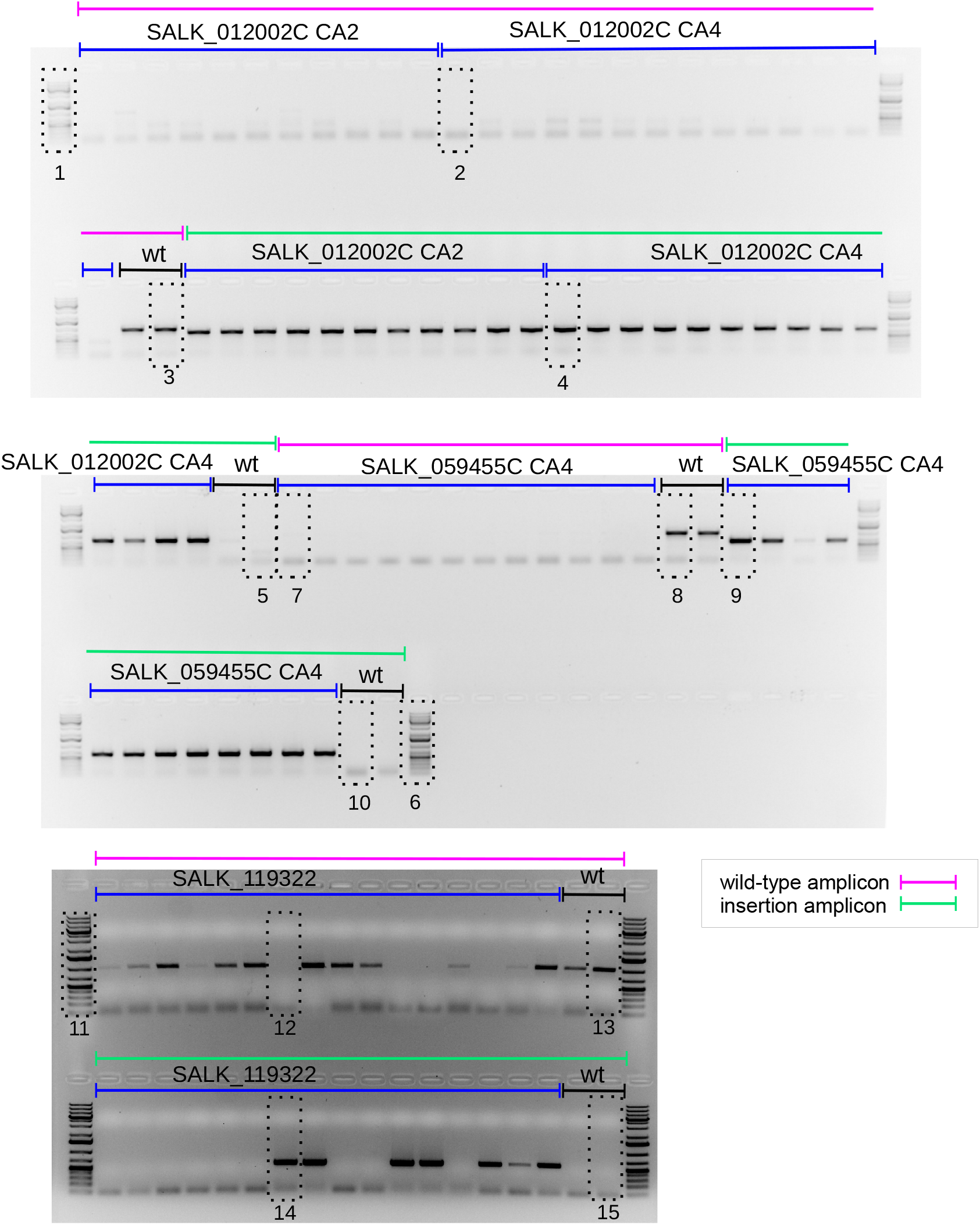
Source data for Figure 2 - Uncropped gel images. PCR products using genomic DNA from individual plants using primers specific to wild-type or insertion sequences. Sections cropped and reassembled for Figure 2 are boxed. Numbers map to position in Figure 2.

**Supplementary Table 1.**
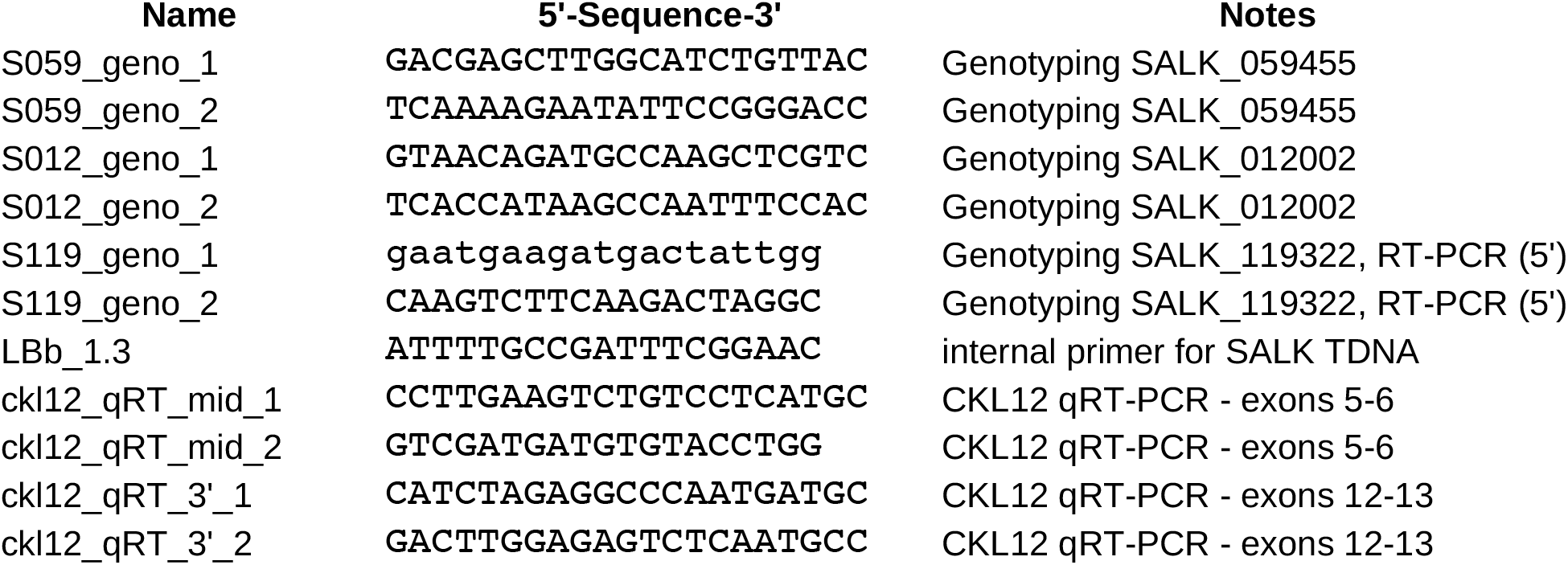
Primer Sequences.

## REFERENCES

1. Peters, J. M., McKay, R. M., McKay, J. P. & Graff, J. M. Casein kinase I transduces Wnt signals. Nature 401, 345–350 (1999).

2. Vielhaber, E., Eide, E., Rivers, A.Gao, Z.-H. & Virshup, D. M. Nuclear Entry of the Circadian Regulator mPER1 Is Controlled by Mammalian Casein Kinase I ε. Mol. Cell. Biol. 20, 4888–4899 (2000).

3. Cheong, J. K. & Virshup, D. M. Casein kinase 1: Complexity in the family. Int. J. Biochem. Cell Biol. 43, 465–469 (2011).

4. Knippschild, U. et al. The CK1 Family: Contribution to Cellular Stress Response and Its Role in Carcinogenesis. Front. Oncol. 4, 96 (2014).

5. Graves, P. R. & Roach, P. J. Role of COOH-terminal Phosphorylation in the Regulation of Casein Kinase Iδ (*). J. Biol. Chem. 270, 21689–21694 (1995).

6. Cegielska, A., Gietzen, K. F., Rivers, A. & Virshup, D. M. Autoinhibition of Casein Kinase I ε (CKIε) Is Relieved by Protein Phosphatases and Limited Proteolysis *. J. Biol. Chem. 273, 1357–1364 (1998).

7. Harold, R. L. et al. Isoform-specific C-terminal phosphorylation drives autoinhibition of Casein kinase 1. Proc. Natl. Acad. Sci. U. S. A. 121, e2415567121.

8. Ono, A. et al. 3, 4-Dibromo-7-Azaindole Modulates Arabidopsis Circadian Clock by Inhibiting Casein Kinase 1 Activity. Plant Cell Physiol. doi:10.1093/pcp/pcz183.

9. Saito, A. N. et al. Structure–function study of a novel inhibitor of the casein kinase 1 family in Arabidopsis thaliana. Plant Direct 3, e00172 (2019).

10. Uehara, T. N. et al. Casein kinase 1 family regulates PRR5 and TOC1 in the Arabidopsis circadian clock. Proc. Natl. Acad. Sci. 201903357 (2019) doi:10.1073/pnas.1903357116.

11. Tan, S.-T., Dai, C.Liu, H.-T. & Xue, H.-W. Arabidopsis Casein Kinase1 Proteins CK1.3 and CK1.4 Phosphorylate Cryptochrome2 to Regulate Blue Light Signaling. Plant Cell Online 25, 2618–2632 (2013).

12. Liu, Q. et al. Molecular basis for blue light-dependent phosphorylation of Arabidopsis cryptochrome 2. Nat. Commun. 8, (2017).

13. Ben-Nissan, G. et al. Arabidopsis Casein Kinase 1-Like 6 Contains a Microtubule-Binding Domain and Affects the Organization of Cortical Microtubules. Plant Physiol. 148, 1897–1907 (2008).

14. Ben-Nissan, G., Yang, Y. & Lee, J.-Y. Partitioning of casein kinase 1-like 6 to late endosome-like vesicles. Protoplasma 240, 45–56 (2010).

15. Cui, Y. et al. Arabidopsis casein kinase 1-like 2 involved in abscisic acid signal transduction pathways. J. Plant Interact. 9, 19–25 (2014).

16. Zhao, X. et al. CKL2 mediates the crosstalk between abscisic acid and brassinosteroid signaling to promote swift growth recovery after stress in Arabidopsis. J. Integr. Plant Biol. n/a, (2022).

17. Li, Y. et al. Promoters of Arabidopsis Casein kinase I-like 2 and 7 confer specific high-temperature response in anther. Plant Mol. Biol. (2018) doi:10.1007/s11103-018-0760-7.

18. Chory, J., Nagpal, P. & Peto, C. A. Phenotypic and Genetic Analysis of det2, a New Mutant That Affects Light-Regulated Seedling Development in Arabidopsis. Plant Cell 3, 445–459 (1991).

19. Li, J., Nagpal, P., Vitart, V., McMorris, T. C. & Chory, J. A role for brassinosteroids in light-dependent development of Arabidopsis. Science 272, 398–401 (1996).

20. Choe, S. et al. The Arabidopsis dwarf1 Mutant Is Defective in the Conversion of 24-Methylenecholesterol to Campesterol in Brassinosteroid Biosynthesis. Plant Physiol. 119, 897–908 (1999).

21. Nagata, N., Min, Y. K., Nakano, T., Asami, T. & Yoshida, S. Treatment of dark-grown Arabidopsis thaliana with a brassinosteroid-biosynthesis inhibitor, brassinazole, induces some characteristics of light-grown plants. Planta 211, 781–790 (2000).

22. Clouse, S. D., Langford, M. & McMorris, T. C. A brassinosteroid-insensitive mutant in Arabidopsis thaliana exhibits multiple defects in growth and development. Plant Physiol. 111, 671–678 (1996).

23. Li, J., Nam, K. H., Vafeados, D. & Chory, J. BIN2, a New Brassinosteroid-Insensitive Locus in Arabidopsis. Plant Physiol. 127, 14–22 (2001).

24. Yin, Y. et al. BES1 Accumulates in the Nucleus in Response to Brassinosteroids to Regulate Gene Expression and Promote Stem Elongation. Cell 109, 181–191 (2002).

25. Wang, Z.-Y. et al. Nuclear-Localized BZR1 Mediates Brassinosteroid-Induced Growth and Feedback Suppression of Brassinosteroid Biosynthesis. Dev. Cell 2, 505–513 (2002).

26. Yin, Y. et al. A New Class of Transcription Factors Mediates Brassinosteroid-Regulated Gene Expression in Arabidopsis. Cell 120, 249–259 (2005).

27. Nemhauser, J. L., Mockler, T. C. & Chory, J. Interdependency of Brassinosteroid and Auxin Signaling in Arabidopsis. PLoS Biol 2, e258 (2004).

28. Bouquin, T., Meier, C., Foster, R., Nielsen, M. E. & Mundy, J. Control of Specific Gene Expression by Gibberellin and Brassinosteroid. Plant Physiol. 127, 450–458 (2001).

29. Neff, M. M. et al. BAS1: A gene regulating brassinosteroid levels and light responsiveness in Arabidopsis. Proc. Natl. Acad. Sci. U. S. A. 96, 15316–15323 (1999).

30. Luccioni, L. G., Oliverio, K. A., Yanovsky, M. J., Boccalandro, H. E. & Casal, J. J. Brassinosteroid Mutants Uncover Fine Tuning of Phytochrome Signaling. Plant Physiol. 128, 173–181 (2002).

31. Sun, Y. et al. Integration of Brassinosteroid Signal Transduction with the Transcription Network for Plant Growth Regulation in Arabidopsis. Dev. Cell 19, 765–777 (2010).

32. Schneider, C. A., Rasband, W. S. & Eliceiri, K. W. NIH Image to ImageJ: 25 years of image analysis. Nat. Methods 9, 671–675 (2012).

33. Kloss, B. et al. The Drosophila Clock Gene double-time Encodes a Protein Closely Related to Human Casein Kinase Iε. Cell 94, 97–107 (1998).

34. Lowrey, P. L. et al. Positional Syntenic Cloning and Functional Characterization of the Mammalian Circadian Mutation tau. Science 288, 483–492 (2000).

35. Preuss, F. et al. Drosophila doubletime Mutations Which either Shorten or Lengthen the Period of Circadian Rhythms Decrease the Protein Kinase Activity of Casein Kinase I. Mol. Cell. Biol. 24, 886–898 (2004).

36. O’Malley, R. C. et al. Cistrome and Epicistrome Features Shape the Regulatory DNA Landscape. Cell 165, 1280–1292 (2016).

37. Pan, X., Li, Y. & Stein, L. Site Preferences of Insertional Mutagenesis Agents in Arabidopsis. Plant Physiol. 137, 168–175 (2005).

38. Baulcombe, D. C., Saunders, G. R., Bevan, M. W., Mayo, M. A. & Harrison, B. D. Expression of biologically active viral satellite RNA from the nuclear genome of transformed plants. Nature 321, 446–449 (1986).

39. Srivastava, A. K., Lu, Y., Zinta, G., Lang, Z. & Zhu, J.-K. UTR dependent control of gene expression in plants. Trends Plant Sci. 23, 248–259 (2018).

